# Accounting for Errors in Low Coverage High-Throughput Sequencing Data when Constructing Genetic Maps using Biparental Outcrossed Populations

**DOI:** 10.1101/249722

**Authors:** Timothy P. Bilton, Matthew R. Schofield, Michael A. Black, David Chagné, Phillip L. Wilcox, Ken G. Dodds

## Abstract

Next generation sequencing is an efficient method that allows for substantially more markers than previous technologies, providing opportunities for building high density genetic linkage maps, which facilitate the development of non-model species’ genomic assemblies and the investigation of their genes. However, constructing genetic maps using data generated via high-throughput sequencing technology (e.g., genotyping-by-sequencing) is complicated by the presence of sequencing errors and genotyping errors resulting from missing parental alleles due to low sequencing depth. If unaccounted for, these errors lead to inflated genetic maps. In addition, map construction in many species is performed using full-sib family populations derived from the outcrossing of two individuals, where unknown parental phase and varying segregation types further complicate construction. We present a new methodology for modeling low coverage sequencing data in the construction of genetic linkage maps using full-sib populations of diploid species, implemented in a package called GUSMap. Our model is based on an extension of the Lander-Green hidden Markov model that accounts for errors present in sequencing data. Results show that GUSMap was able to give accurate estimates of the recombination fractions and overall map distance, while most existing mapping packages produced inflated genetic maps in the presence of errors. Our results demonstrate the feasibility of using low coverage sequencing data to produce genetic maps without requiring extensive filtering of potentially erroneous genotypes, provided that the associated errors are correctly accounted for in the model.

---

The emergence of high-throughput sequencing methods that multiplex large numbers of individuals has provided a cost-effective approach to perform genome-wide genotyping and discovery of genetic variation. Two of the primary multiplexing sequencing methods are whole genome sequencing, and reduced representation approaches, including whole-exome sequencing (Hodges *et al*. 2007), restriction-site associated DNA sequencing (Baird *et al*. 2008), and genotyping-by-sequencing (Elshire *et al*. 2011) among others (Heffelfinger *et al*. 2014). The introduction of these methods has led to the rapid increase in both the number of species being sequenced, especially non-model species (Ellegren 2014), and the number of markers available for analysis. Consequently, these methods provide opportunities to construct more dense genetic linkage maps compared with previous technologies, which is particularly useful in scenarios where alternative high density marker systems are infeasible (expensive to establish and validate). Genetic maps are important as they facilitate the investigation of many species in terms of their genes, such as associating phenotypes to the genome via quantitative trait loci, validating assemblies, ordering contigs in assemblies, and performing comparative genome analyses (Cheema and Dicks 2009; Liu *et al*. 2014).

Constructing linkage maps using sequencing data is complicated by the presence of two types of missing data that can result when the sequencing depth is low. The first is a missing genotype resulting from no alleles being called, while the second consists of a heterozygous genotype being called as homozygous due to only one of the parental alleles being sequenced at a particular locus (Fragoso *et al*. 2016; Dodds *et al*. 2015). The latter type is particularly problematic as it usually behaves like a genotyping error, which increases the frequency of inferred recombinations and results in inflated linkage maps (Cartwright *et al*. 2007; Lincoln and Lander 1992; Cheema and Dicks 2009). Typically, genotyping errors resulting from low sequencing coverage are removed via filtering, such as setting genotypes with an associated read depth below some threshold value to missing (Gardner *et al*. 2014; Mousavi *et al*. 2016) or using genotype quality scores to discard uncertain genotype calls (Mousavi *et al*. 2016; Hyma *et al*. 2015; Chen *et al*. 2014). Nevertheless, this requires sequencing at a higher depth, which results in fewer individuals being sequenced and fewer utilized loci for a given cost, and can leave a large proportion of the original data unused. Several algorithms have been developed for imputing missing genotypes and correcting erroneous genotypes in low coverage genome sequencing data (Fragoso *et al*. 2016; Swarts *et al*. 2014; Huang *et al*. 2014; Spindel *et al*. 2013), however, all of these algorithms are designed only for inbred populations and are not applicable to outcrossed full-sibling (full-sib) families. Recently, two software packages have been developed for performing linkage mapping in full-sib families using sequencing data. These are Lep-MAP (Rastas *et al*. 2013, 2016) and HighMap (Liu *et al*. 2014), both of which address the computational problem associated with high density maps but are not specifically designed to handle low coverage sequencing data.

Another complication is the presence of sequencing errors, reads where the base has been called incorrectly. In contrast to errors caused by low read depth, sequencing errors can result in homozygotes being called as heterozygotes. Nevertheless, both types of errors lead to inflated genetic distances if not taken into account. One approach for removing sequencing errors involves detecting double recombinants at very short distances and either correcting the genotypes (e.g., a double recombinant becomes nonrecombinant) or setting the genotypes resulting in double recombinants as missing (Cheema and Dicks 2009; Liu *et al*. 2014; Van Os *et al*. 2005; Wu *et al*. 2008). The problem with correcting double recombinants is the possibility of false positives, particularly if the chromosomal order is inaccurate (Wu *et al*. 2008), while erroneous genotype calls on the outside loci cannot be detected using this approach. An alternative is to account for these errors by including additional parameters in the model (Cartwright *et al*. 2007; Rastas *et al*. 2013), although estimation of these parameters is not always straightforward when the error rate is unknown.

Linkage mapping in plants has often been applied to inbred populations derived from the cross of two fully homozygous parents (e.g., recombinant inbred lines, double haploids) (Maliepaard *et al*. 1997; Grattapaglia and Sederoff 1994), although this is dependent upon the breeding system of the species. For many plant species and most animals species, self-incompatible, severe inbreeding depression or long generation times prevent the production of inbred lines, where an alternative mapping population commonly used is an outbred full-sib family derived from the crossing of two unrelated individuals (Schneider 2005; Singh and Singh 2015). Examples where outbred populations have been particularly utilized in linkage mapping range from forest trees to forages (Grattapaglia and Sederoff 1994; Devey *et al*. 1994; Plomion *et al*. 1995; Wilcox *et al*. 2001; Butcher *et al*. 2002; Faville *et al*. 2004; Griffiths *et al*. 2013). However, building linkage maps in outcrossed populations is complicated by loci having different segregation types (e.g., the number of alleles segregating in each parent) and unknown parental phase (Maliepaard *et al*. 1997; Lu *et al*. 2004). An early approach for performing linkage mapping in these populations was the pseudo-testcross strategy (Grattapaglia and Sederoff 1994), which maps the paternal and maternal meioses independently. However, when there are loci segregating in both parents, this approach does not use all available information, while integration of the two parental maps is complicated (Maliepaard *et al*. 1997; Van Ooijen 2011). An alternative approach is to model both meioses simultaneously using a multipoint likelihood model, provided there is a sufficient number of loci segregating in both parents (Van Ooijen 2011). One such model is the Lander-Green hidden Markov model (HMM) for general pedigrees (Lander and Green 1987). Although applicable to full-sib family populations, computation of the Lander-Green HMM is infeasible for moderate-to-large pedigrees (Thompson 2000). Several variants of this model derived specifically for full-sib family populations in diploid species have been suggested (Ling 2000; Wu *et al*. 2002; Tong *et al*. 2010), which reduces the computational complexity by exploiting the conditional independence between individuals given the parental phase.

In this article, we describe a new statistical method that adjusts for bias in map length estimation due to errors in genotypic data derived from sequencing. Our method is based on the Lander-Green HMM for full-sib families in diploid species (Ling 2000; Tong *et al*. 2010; Wu *et al*. 2002) that is applicable to multi-family and sex-specific situations, but includes an additional component to account for errors associated with sequencing data. The performance of the methodology presented here is tested and compared with existing full-sib family software packages using simulations and a real sequencing data set.

## Materials and Methods

### Segregation type and parental phase

In full-sib families, the combination of alleles found in the parental genotypes, referred to as segregation type (ST), varies from locus to locus. A complete classification of all STs in a full-sib family for diploid species has been given by Maliepaard *et al*. (1997). For sequencing data in diploid species that typically consists only of single nucleotide polymorphisms (SNPs), the number of different alleles found at a given locus in the parents is usually two. Consequently, the relevant STs are *AB × AB, AB × AA*, *AB × BB*, *AA × AB* and *BB × AB*, where *A* denotes the reference allele, *B* denotes the alternate allele (maternal × paternal), and *AA*, *AB* and *BB* denote the reference homozygous, heterozygous and alternate homozygous genotypes, respectively. Using the standard nomenclature of Groover *et al*. (1994), we refer to the STs *AB × AB* as both-informative (BI), *AB × AA* and *AB × BB* as maternal-informative (MI), and *AA × AB* and *BB × AB* as paternal-informative PI_*B*_. To distinguish between the two MI and PI STs, we refer to *AB × AA* as MI_*A*_, *AB × BB* as MI_*B*_, *AA × AB* as PI_*A*_ and *BB × AB* as PI_*B*_. The STs of *AA × AA, AA × BB, BB × AA* and BB × BB are also possible, although they are not usually classified as they provide no information of recombination in either parent, and are hence referred to as uninformative (U). Uninformative loci are included in this classification since it is possible for a locus to be uninformative in one family but informative in another.

We let 
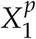
 denote the allele on the paternally derived chromosome of the paternal parent, 
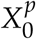
 denote the allele on the maternally derived chromosome of the paternal parent, 
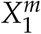
 denote the allele on the paternally derived chromosome of the maternal parent, and 
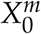
 denote the allele on the maternally derived chromosome of the maternal parent. The ordered parental genotype pair (OPGP) is defined as the unique combination of 
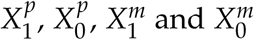
. Across the four STs of BI, PI, MI and U, there are sixteen distinct OPGPs (Table 1). Specification of the OPGP for all loci is equivalent to determining the parental haplotypes and consequently the allelic phase of the parents.

**Table 1.** The OPGPs for both-informative, paternal-informative, maternal-informative and uninformative biallelic loci in a full-sib family

| ST | OPGP | $X_1^p$ | $X_0^p$ | $X_1^m$ | $X_0^m$ |
| --- | --- | --- | --- | --- | --- |
| BI | 1 | $A$ | $B$ | $A$ | $B$ |
| | 2 | $B$ | $A$ | $A$ | $B$ |
| | 3 | $A$ | $B$ | $B$ | $A$ |
| | 4 | $B$ | $A$ | $B$ | $A$ |
| $PI_A$ | 5 | $A$ | $B$ | $A$ | $A$ |
| | 6 | $B$ | $A$ | $A$ | $A$ |
| $PI_B$ | 7 | $A$ | $B$ | $B$ | $B$ |
| | 8 | $B$ | $A$ | $B$ | $B$ |
| $MI_A$ | 9 | $A$ | $A$ | $A$ | $B$ |
| | 10 | $A$ | $A$ | $B$ | $A$ |
| $MI_B$ | 11 | $B$ | $B$ | $A$ | $B$ |
| | 12 | $B$ | $B$ | $B$ | $A$ |
| U | 13 | $A$ | $A$ | $A$ | $A$ |
| | 14 | $A$ | $A$ | $B$ | $B$ |
| | 15 | $B$ | $B$ | $A$ | $A$ |
| | 16 | $B$ | $B$ | $B$ | $B$ |

### Data and models

We begin by assuming that there are no errors present. In such a case, the data are denoted by G_*fij*_, the true genotype call (*AA, AB* or *BB*) for individual *i* in family *f* at locus *j* for *f* = 1, …, *F*, *i* = 1, …, N_*f*_ and *j* = 1, …, *M*, where *F* is the total number of families, N_*f*_ is the number of individuals in family *f* and *M* is the total number of loci. We denote the vector (length N) of true genotypes at locus *j* by 
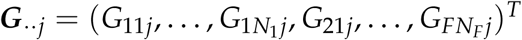
, where 
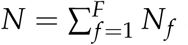
 and the superscript *T* denotes the transpose. The latent inheritance vectors are denoted **S**_*fij*_ = (s_1_, s_2_)^*T*^, where *s*_1_ is the inheritance from the paternal parent and *s*_2_ is the inheritance from the maternal parent. The value of *s*_k_ is 0 if the allele is derived from the parent’s maternal chromosome and 1 if the allele is derived from the parent’s paternal chromosome for *k* = 1, 2. We denote the inheritance vector (length 2*N*) for all individuals at locus *j* by 
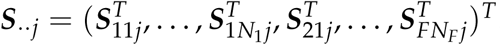

***Lander-Green HMM:*** For multilocus analysis in general pedigrees, Lander and Green (1987) proposed using the HMM

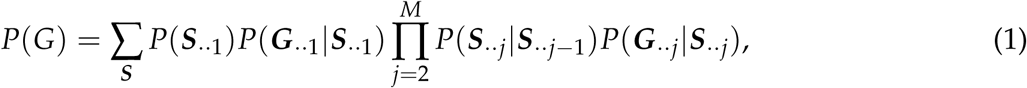

where 
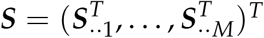
. In HMM theory 
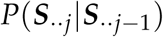
 is known as the transmission probability, 
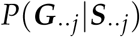
 as the emission probability, and P(**S**_..1_) as the initial distribution. Usually, the initial distribution is taken to be 2*N* independent Bernoulli trials (Ling 2000; Tong *et al*. 2010).

***HMM for full-sib families:*** In its original form, the Lander-Green HMM likelihood can be computed in *O(N^2^M)* time using the forward-backward algorithm of Baum *et al*. (1970). Computing the Lander-Green HMM likelihood quickly becomes infeasible for pedigrees of moderate-to-large sizes. In full-sib family populations, individuals within and between families are conditionally independent given the OPGPs (e.g., parental phases). If z_*fj*_ denotes the OPGP at locus *j* in family *f*, then the HMM for full-sib family populations can be expressed as,

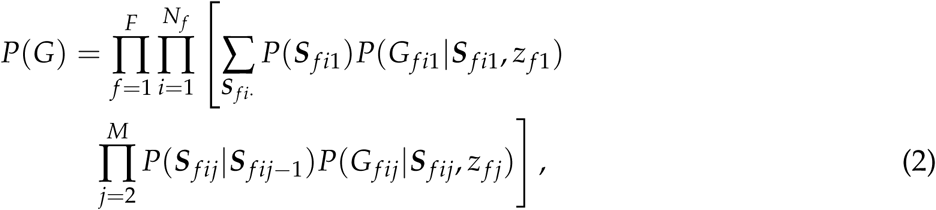

where 
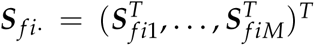
. Using model (2), the computational time is reduced to *O(NM)*, provided that the OPGPs are known for all families.

For a single individual in a full-sib family, the four inheritance vectors, **S**_
*fij*_, that are possible in the HMM are (0, 0), (0, 1), (1, 0) and (1, 1). Let 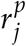
 and 
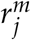
 denote the paternal and maternal recombination fraction respectively between locus *j* and locus *j* + 1, where these recombination fractions are constrained to the interval [0, 0.5]. For model (2), the transition probabilities, 
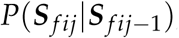
, are given in Table 2, while the emission probabilities, 
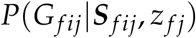
, are given in Table S1 in File S1. When the genotype is missing, the emission probability is 
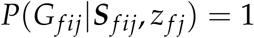
 for all inheritance vectors.

**Table 2.** Transmission probabilities for the full-sib family HMM.

| $S_{fij-1}$ | $S_{fij}$ | | | |
| --- | --- | --- | --- | --- |
|  | (0,0) | (0,1) | (1,0) | (1,1) |
| (0,0) | $(1 - r_j^p)(1 - r_j^m)$ | $(1 - r_j^p)r_j^m$ | $r_j^p(1 - r_j^m)$ | $r_j^p r_j^m$ |
| (0,1) | $(1 - r_j^p)r_j^m$ | $(1 - r_j^p)(1 - r_j^m)$ | $r_j^p r_j^m$ | $r_j^p(1 - r_j^m)$ |
| (1,0) | $r_j^p(1 - r_j^m)$ | $r_j^p r_j^m$ | $(1 - r_j^p)(1 - r_j^m)$ | $(1 - r_j^p)r_j^m$ |
| (1,1) | $r_j^p r_j^m$ | $r_j^p(1 - r_j^m)$ | $(1 - r_j^p)r_j^m$ | $(1 - r_j^p)(1 - r_j^m)$ |

To compute the likelihood of the full-sib family HMM, forward recursion is used. Define *α_fij_*(*S_fij_*) as the forward probability which satisfies the relations

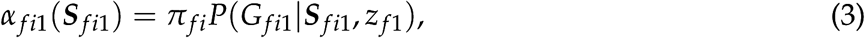

where *π_fi_* = P(***S**_fi1_*), and

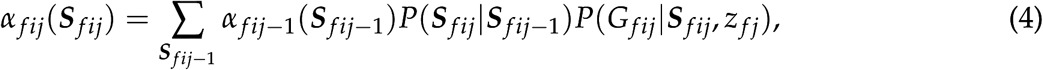

for *j* = 2, …, *M*. Under the assumption that the initial distribution is 2*N* independent Bernoulli trials, π_*fi*_ = 1/4 for all *f,i*. The likelihood of the HMM for individual *i* in family *f* is

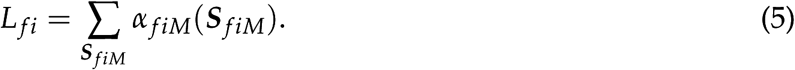

As individuals within and between families are conditionally independent given the OPGPs of all the parents, the likelihood for multiple full-sib families is

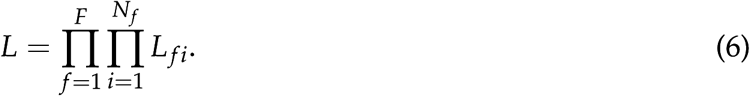

In situations where some loci are uninformative in the maternal or paternal parent across all families, a slight adjustment to the parametrization of the model is required. If the paternal (maternal) genotype at locus *j* is homozygous in every family or the paternal (maternal) genotypes at all loci from locus 1 to *j* − 1 are homozygous in every family, then the recombination fraction 
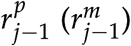
 cannot be estimated and therefore is excluded from the model. Under this parametrization, the sex-specific recombination fraction 
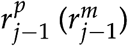
 is now interpreted as the probability of a recombination in the paternal (maternal) parent between locus *j* and the previous locus which is segregating in the paternal (maternal) parent. When the sex-specific recombination fractions are assumed equal 
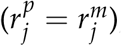
 this adjustment to the parametrization is not required.

***Incorporating errors in the Lander-Green HMM:***When there is error present in the sequencing data, the genotypes, G_*fij*_, are latent. The observed data are the number of reads for the reference allele, *A*, and alternate allele, *B*. We denote the number of reads for the reference allele observed for individual *i* in family f at locus *j* 
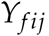, where 
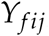 is an integer value between 0 and *d*_*fij*_, and *d*_*fij*_ is the sequencing depth at locus *j* for individual *i* in family *f*. The sequencing depth, *d*_*fij*_, is equal to the sum of the number of reads for the reference and alternate alleles. We denote the vector (length N) of reference allele counts at locus *j* by 
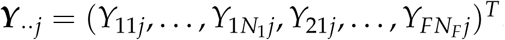
 If 
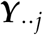 is conditionally independent between loci given ***G**_¨j_*, then the extended HMM for sequencing data becomes

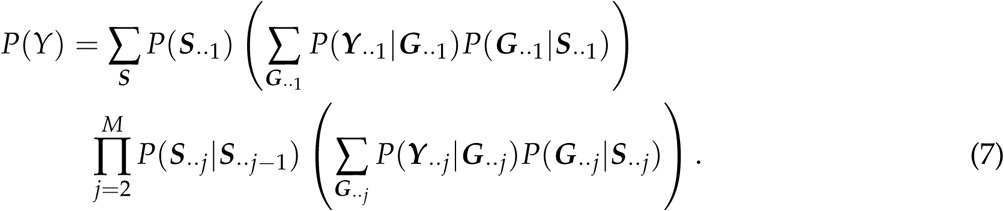

The transmission probabilities in model (7) are the same as in model (1). The emission probability is 
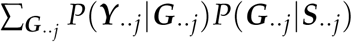
 conditional on the sequencing depth *d*_*fij*
_.

*Full-sib HMM for sequencing data:* If the number of reference alleles observed in the sequencing data, 
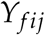, is conditionally independent between individuals given the true genotypes, G_*fij*_, then the full-sib family HMM for sequencing data is

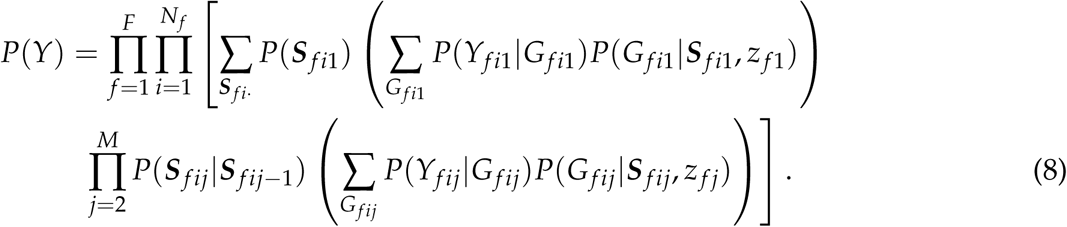

The only change to model (8) compared with model (2) is in the emission probabilities, which requires specifying the conditional probabilities 
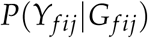
 Suppose that Y_fij_ arises from a random binomial sample of the alleles found in G_*fij*_ (Dodds *et al*. 2015) and suppose that sequencing errors occur independently between reads, then

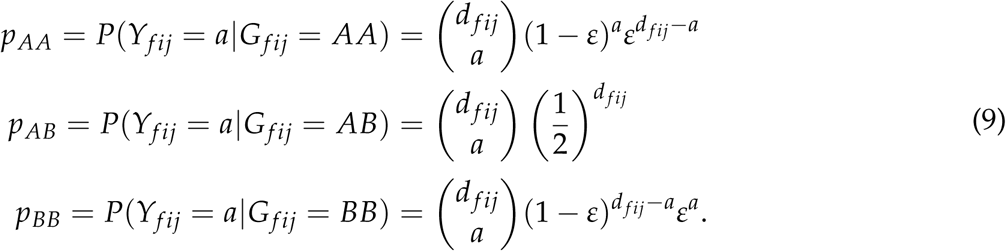

See File S2 for derivation of these probabilities. Under these assumptions, the emission probabilities, 
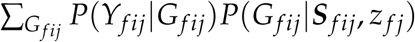
 for model (8) can be derived (Table S2 in File S1). Consequently, the likelihood of the HMM for sequencing data equates to Eq (6) with the emission probability 
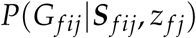
 replaced by 
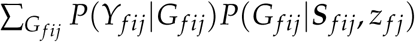
.

### Inferring OPGPs

The likelihoods for full-sib families derived in the previous sections assume that the OPGPs (or parental phases) are known. In practice, this information is unknown, although the OPGPs can, in some cases, be inferred from the grandparental genotype information. Nevertheless, if there is no grandparental information, then inference of the OPGPs using progeny genotypes (assuming parents are known and accurately genotyped) is required.

We initialize the value of *z*_*fj*_ for each locus to a default value, that is, we initialize *z*_*fj*_ = 1 if the locus is BI, *z*_*fj*_ = 5 if the locus is PI_*A*_, z _fj_ = 7 if the locus is PI_*B*_, *z*_*fj*_ = 9 if the locus is MI_*A*_, and *z*_*fj*_ = 11 if the locus is MI_*B*_. Inference of the OPGPs for family *f* can be achieved by relaxing the constraint on 
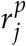
 and 
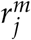
 such that 
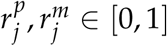
 and maximising the likelihood

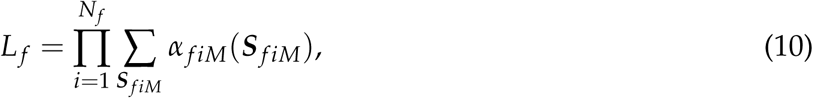

where *α*_*fiM*_(***S***_
*fiM*_) is defined as in Eqs (3) and (4), and the emission probability is 
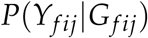
 for inference under model (2) but 
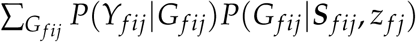
 for inference under model (8). The OPGP of locus *j* = 2, …, *M* can be inferred relative to the previous OPGPs based on whether the maximum likelihood estimates of 
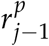
 and/or 
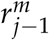
 are greater than or less than 0.5, where the OPGP for the first locus is set to a baseline value depending on its ST (see File S2 for details).

### Implementation

An implementation of the new methodology presented in this paper can be found in the GUSMap (Genotyping Uncertainty with Sequencing data and linkage MAPping) software, which is freely available as a package for the programming language R (R Core Team 2017) and can be downloaded from https://github.com/tpbilton/GUSMap. In this package, numerical maximization methods are used to compute the maximum likelihood estimates of the likelihoods. Specifically, we use the ‘BFGS’ method implemented in the **optim()** function with the likelihood functions written in C to reduce computational time. Since an unconstrained numeric optimizer is used, the likelihoods are solved with respect to transformed model parameters using the transformation

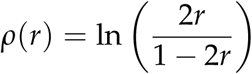

for the recombination fractions in all the likelihoods except for likelihood (10) where the logit transformation is used, and the logit transformation for the sequencing error parameter, *ε*. The maximum likelihood estimates for the parameters are computed by back transforming the transformed parameter estimates.

### Software comparison

Using simulated and real data, the performance of GUSMap v0.1.0 (GM) was compared to the four linkage mapping software packages: CRI-MAP 2.507 (CM) (Green *et al*. 1990), JoinMap 4.1 (JM) (Van Ooijen 2011), Lep-MAP2 (Rastas *et al*. 2016) and OneMap v2.0-4 (OM) (Margarido *et al*. 2007), all of which are commonly used for full-sib family populations. In general, the default parameter settings were used, except for JM and Lep-MAP2. In JM, the threshold for determining linkage was set to zero in order for a complete map to be computed in every data set and the maximum likelihood algorithm (Van Ooijen 2011) was used. For Lep-MAP2, detection of duplicate loci was removed (argument removeDuplicates=0). In addition, the methodology of Lep-MAP2 includes estimation of an error parameter for each locus and was implemented using two sets of parameter options. The first corresponds to the model that includes the error parameters, referred to as LM2*ε*, while the second corresponds to the exclusion of all error parameters (using arguments learnErrorParameters=0 and initError=0) and is referred to as LM2.

With sequencing data, some genotype calls may result in apparent Mendelian errors, which occur when a genotype call for a PI_*A*_ or MI_*A*_ locus is homozygous for the alternate allele, or a genotype call for a PI_*B*_ or MI_*B*_ locus is homozygous for the reference allele. Genotype calls determined to be a Mendelian error were set as heterozygous, since most of the standard linkage mapping packages cannot handle data sets with these errors present. Mendelian errors were not corrected with GM as they are accounted for in the HMM. In addition, some heterozygous genotype calls in the sequencing data were supported by more than nine reads for one allele but only a single read for the other allele. As these genotype calls are likely to be sequencing errors, they were set to missing for the standard packages, but not for GM, as they provide information used to estimate the sequencing error parameter *ε*.

### Simulation

Sequencing data were simulated using the following procedure. Inheritance vectors for progeny were generated based on the true parental recombination values assuming no interference and equal probability of the first locus being derived from either parent. These inheritance vectors were converted to genotype calls for a pre-specified set of OPGPs. From these true genotype calls, the simulated sequencing data sets were generated as follows:

- A sequencing depth at each locus in each individual was generated by simulating realizations from a negative binomial distribution with mean 
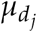
 and dispersion parameter of 2, that is

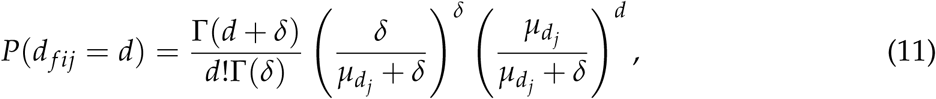

where *δ* = 2, 
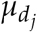
 corresponds to the mean sequencing depth for locus *j* and Γ(.) denotes the gamma function.

- A sample of *d*_*fij*_ alleles are found by randomly sampling the alleles of the true genotype, G_*fij*_, with replacement, where a miscall of the sampled allele (e.g., a *B* allele called as *A* and vice versa) occurred with probability *ε*.

Two sets of simulations were conducted. In the first set, the performance of the five software packages is examined and compared under different read mean depths and sequencing error rates. This set of simulations consisted of simulating a 1,000 single full-sib families (*F* = 1) with a hundred offspring (*N*_1_ = 100), twelve loci (*M = 12*) and a fixed recombination rate of 1% in both parents 
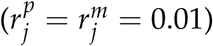
, where the segregation types and OPGPs of the loci are given in Table 3. Different combinations of mean read depth, 
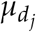
, and sequencing error, *ε*, were used, where the mean readdepth was either low 
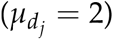
 moderate 
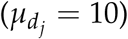
 or high 
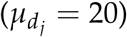
 and the sequencing error rate was either absent (*ε* = 0), small (*ε* = 0.002) or large (*ε* = 0.01). To remove errors associated with low sequencing depth, the simulated data were filtered such that all genotype calls with an associated read depth below some threshold were set to missing. The threshold used was eleven for the high depth setting 
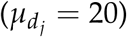
, six for the moderate depth setting 
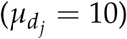
, but wasn’t applied for the low depth setting 
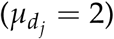
 since an insufficient number of non-missing genotypes would remain. This filtering step was also not performed for GM as it models under-called heterozygous genotypes.

**Table 3.**
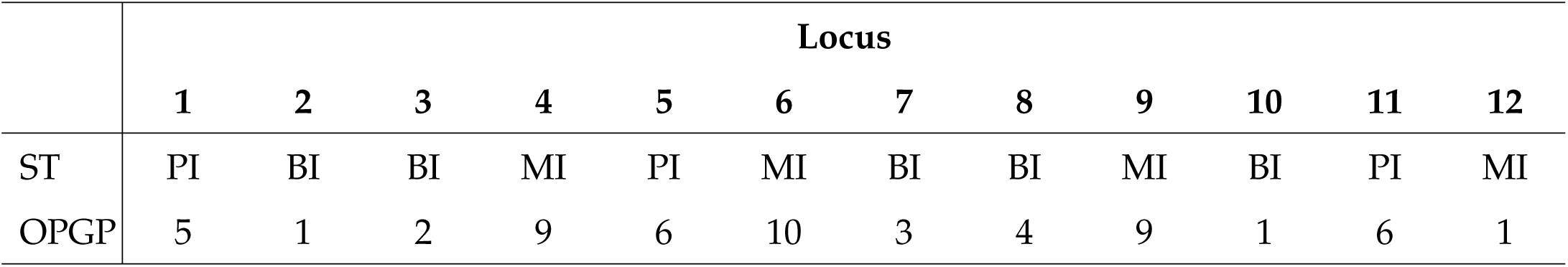
Segregation type and OPGP for loci used in the first set of simulations.

The second set of simulations investigates the optimal sequencing depth for a given sequencing effort (defined as the number of individuals times the number of loci times the mean read depth). The parameters used in this set corresponded with the previous set, with the exception that the sequencing error rate was fixed at 0.2% (*ε* = 0.002), the number of individuals was varied and the mean read depth was set such that an average sequencing effort of 10,000 was maintained. Recombination fractions were estimated using GM assuming a known OPGP.

Code for implementing the simulations is found in File S3. GM, CM, LM2, LM2 *ε*, and OM were all run on a Linux desktop computer with four Intel Core i7-870 central processing units running at 2.93 GHz frequency, while JM was run on a Windows 10 Enterprise desktop computer with four Intel Core i7-3770 central processing units running at 3.40 GHz frequency. As JM has no scripting functionality, it was automated using a custom C# script coupled with a coded user interface test, a program which can automate mouse clicks and keyboard strokes on Windows operating systems.

### Mānuka data

A single full-sib bi-parental family (n = 180) of mānuka (*Leptospermum scoparium*, J.R. Forst. et G. Forst.; Myrtaceae) derived from a reciprocal pair-cross of heterozygous individuals, was genotyped along with the parents, using a genotyping-by-sequencing (GBS) approach (Elshire *et al*. 2011). Samples consisted of young expanding leaves collected from three month old seedlings grown in the glasshouse. Two GBS libraries were prepared based on the Elshire method (Elshire *et al*. 2011) using a double digest with the restriction enzymes A*pe*KI/M*sp*I and sequenced at AgResearch, Invermay, Animal Genomics laboratory. A size selection step was performed on the DNA such that the genomic part of each read was between 27 and 377 base pairs. The samples were sequenced on an Illumina HiSeq 2500 v4 chemistry producing 1x100 single end reads. Each GBS library was sequenced on two lanes of a flow cell generating approximately 29.2 giga base pairs of raw sequence data per lane. The two parents were run on both lanes to obtain higher sequencing depths, while each progeny was run on one of the two lanes. Quality control was performed using DECONVQC (https://github.com/AgResearch/DECONVQC) and KGD (Dodds *et al*. 2015). Three progeny were excluded from further analysis, one due to having a sample call rate below 0.05 while two other samples were identified as being a duplicate of another sample. Sequence reads were mapped using Bowtie2 version 2.1.0 (Langmead and Salzberg 2012) and SNP variants were called using Tassel3 version 3.0.173 (Bradbury *et al*. 2007).

To compare the performance of GM relative to the other packages, only variants called on chromosome 11 were retained for further analysis, with additional filtering performed as follows. SNPs with a minor allele frequency less than 0.05 or 20% or more missing genotypes were discarded. The ST of each SNP was inferred based on the parental genotypes provided that the read depth for both parents was greater than five, where SNPs were discarded if the ST could not be inferred. A segregation test was performed on each SNP using a chi-square test, where a P-value of 0.05 was used and the expected counts were adjusted for low read depth calls (see File S2 for details). To ensure that each tag was only represented by a single variant, adjacent SNPs were placed into bins if the distance separating them was less than 180 base pairs, with one SNP from each bin retained for the final analysis by random selection. After filtering, 680 SNPs remained with 270 PI, 294 MI and 116 BI loci. This data is available in the R package GUSMap.

To assess the ordering of the SNPs called on chromosome 11, heatmaps of the 2-point recombination fraction estimates between all the SNPs segregating in the same parent were produced. GM was used to compute the 2-point recombination fractions (with *ε* = 0), where the phase between the SNP pair was taken as the one which maximized the likelihood value. Linkage maps were computed using two independent sets of SNPs: a low depth set consisting of all the SNPs with a mean read depth below six and a high depth set which was obtained by setting all genotype calls with a read depth below twenty to missing and selecting all the SNPs such that a call rate of at least 80% was maintained. In total, there were 95 Low Depth SNPs and 54 High Depth SNPs.

### Data availability

The mānuka data set used to construct linkage maps is available in the software package GUSMap (https://github.com/tpbilton/GUSMap). Code for generating the simulated data is found in File S3. Supplementary tables and figures are given in File S1 and supplementary methods used in this paper are given in File S2.

## Results

### Simulations

The distribution of the overall map distance estimates obtained using the various software packages in the first set of simulations is given in Figure 1, while the distribution of the recombination fraction estimates for each simulation are given in Figures S1-S9 in File S1. Across all of the simulations, LM2, OM and CM performed similarly and at high depth with no sequencing error, gave relatively unbiased estimates. However, at moderate read depth with no sequencing error, the overall map distance estimates from LM2, OM and CM were slightly larger that the true value, which suggests that the cut-out of six has not removed all the errors associated with low sequencing depth. The bias in the map distance and recombination fractions estimates increased as the level of sequencing error increased for the high and moderate depth scenarios. In comparison to LM2, OM and CM, JM produced maps that were on average slightly longer and more biased across all the low and moderate depth simulations. These inflated maps seem to be driven by biases in the recombination fraction estimates for *r*_4_ and *r*_5_ (see Figures S1-S6 in File S1), particularly when the sequencing error was small or absent. These recombination fraction parameters all include one of the loci in the region between locus 4 and locus 6, where a PI locus is wedged between two MI loci. As JM uses only a 3-point approach, the lack of informativeness between adjacent loci in this region may explain the observed bias.

**Figure 1.**
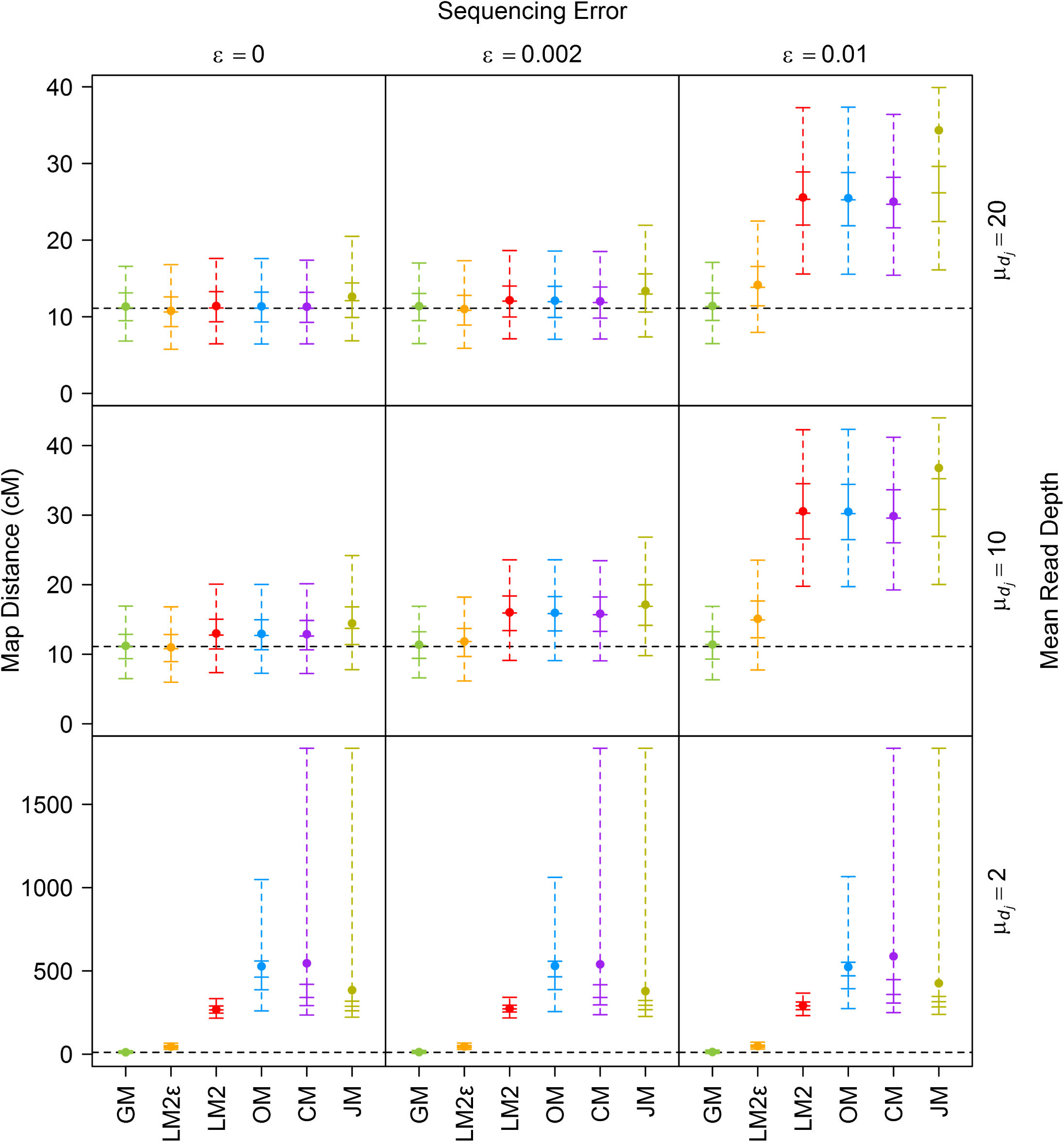
Distribution of the map distance estimates for the first set of simulations across varying mean read depths (rows) and varying sequencing error rates (columns). The solid point represents the mean, the vertical solid line represents the interquartile range, the vertical dashed line represents the range between the 2.5^th^ and 97.5^th^ percentiles, the five horizontal solid lines represent, in ascending order, the 2.5^th^ percentile, lower quantile, median, upper quantile and 97.5^th^ percentile, and the horizontal black dotted line represents the true parameter value. Map distances are in centimorgans (cM) and were computed using the Haldane mapping function.

LM2*ε* was able to produce accurate estimates of the recombination fractions and overall map distance when the sequencing error was absent or low for the moderate and high read depth scenarios. Nevertheless, when the sequencing error was large, LM2*ε* produced biased estimates of the the overall map distance, which was driven by large biases in the recombination fraction estimates that include an outside locus (see Figure S3 and Figure S6 in File S1). At low depth, the four existing software packages all gave very poor map distance estimates across all of the various sequencing error rates, which was expected given the large number of errors in the data sets. Of these methods, LM2*ε* performed the best although its map distance estimates were still approximately four to five times larger than the true value (Figure S10 in File S1). In addition, the recombination fraction estimates for LM2*ε* at low depth were biased (see Figures S7-S9 in File S1), although for the middle sections of the map, the bias was in both directions resulting in less inflation of the overall map distance but a distortion of the distribution of the SNPs across the linkage map. In contrast, GM was the only package which was able to give accurate estimates of the overall map distance and recombination fractions across all the simulation scenarios.

The distribution of the sequencing error estimates obtained from GM are given in Figure 2. For the high and moderate depth simulations, the estimates were relatively accurate, while there was a small bias for the low depth simulations. The variability of the estimates increased as the mean read depth decreased, which is not surprising given that there is more variability in the data at low sequencing depths.

**Figure 2.**
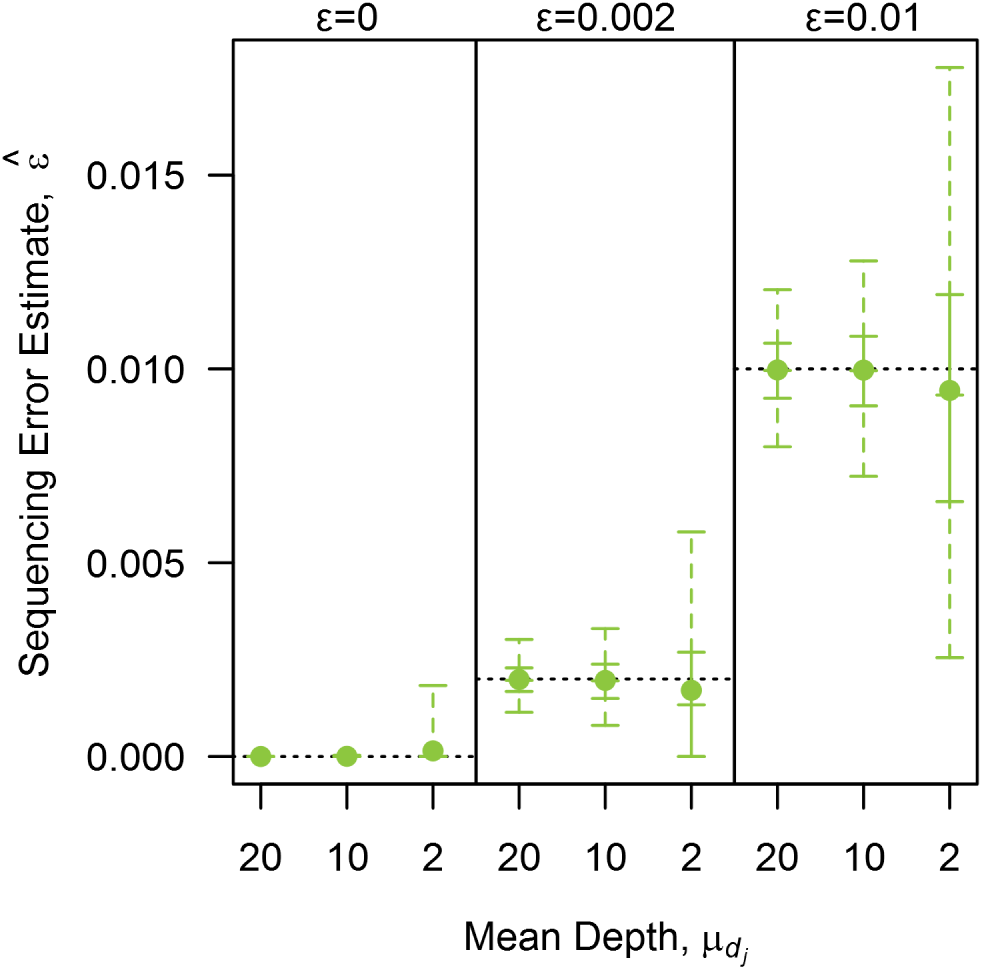
Distribution of the sequencing error estimates obtained from GM for the various combinations of mean read depths and sequencing error rates. The solid point represents the mean, the vertical solid line represents the interquartile range, the vertical dashed line represents the range between the 2.5^th^ and 97.5^th^ percentiles, the five horizontal solid lines represent, in ascending order, the 2.5^th^ percentile, lower quantile, median, upper quantile and 97.5^th^ percentile, and the horizontal black dotted lines represent the true parameter values.

Figure 3 gives the distribution of the computation time required for each package across all of the first set of simulations. Of all the packages, LM2 was the fastest, regardless of whether the error parameters were included, while CM, GM and OM was approximately three times, five and a half times and forty five times slower than LM2 respectively. As JM is a non-scripting program, providing a sensible measure of computation time is difficult. For these simulations, the time recorded was only for the step to compute the map, which on average required four times more computational time than LM2, but did not include the extensive user interaction time needed to import the data and create the required nodes.

**Figure 3.**
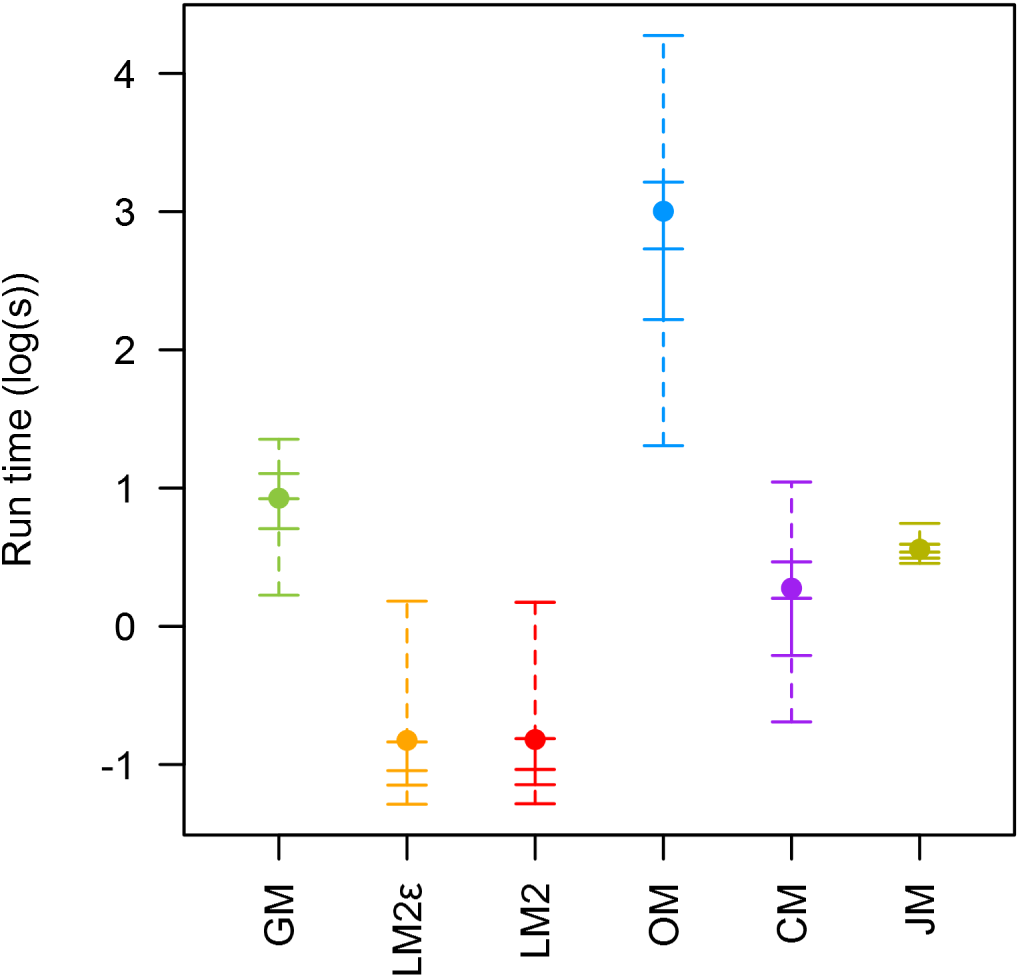
Distribution of the log transformed computational time used on each data set across all nine simulation scenarios for the first set of simulations and for each software package. The solid point represents the mean, the vertical solid line represents the interquartile range, the vertical dashed line represents the range between the 2.5^th^ and 97.5^th^ percentiles, the five horizontal solid lines represent, in ascending order, the 2.5^th^ percentile, lower quantile, median, upper quantile and 97.5^th^ percentile.

The percentage of data sets in which the vector of OPGPs was correctly inferred in the first set of simulations is displayed in Table 4. For the moderate and high depth simulations, all the packages apart from JM were able to correctly infer the parental phase, regardless of the amount of sequencing error present. In contrast, for the low depth simulations, only GM and LM2*ε* were able to correctly infer phase across all the simulations, while OM rarely inferred phase correctly and LM2 incorrectly inferred phase for a few data sets. There were a small number of phasing errors for JM across the various scenarios, with the frequency of these errors increasing as the number of erroneous genotypes increased. For CM, phase inference was not required since it is an implementation of the Lander-Green HMM for general pedigrees.

**Table 4.**
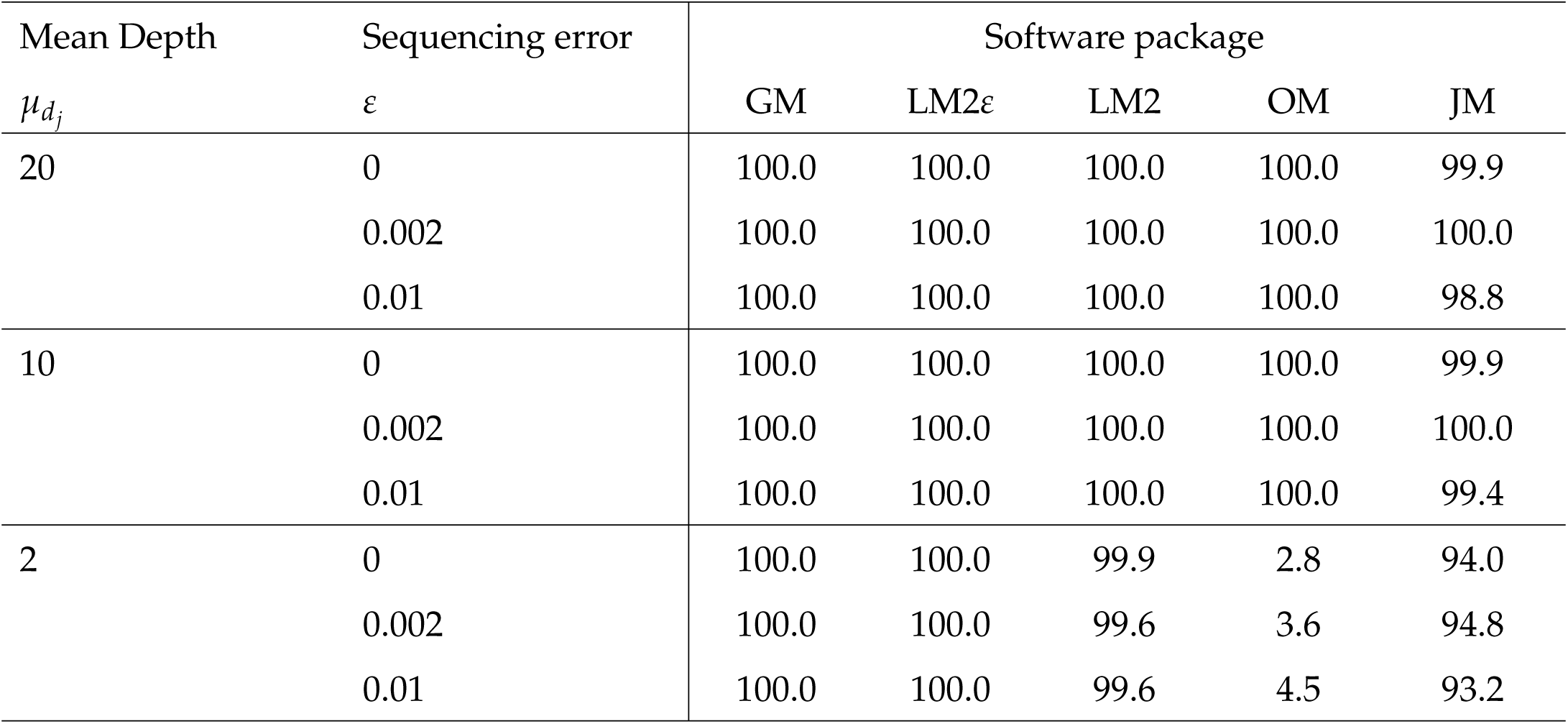
Percentage of simulated data sets in which the vector of OPGPs was correctly inferred

For the second set of simulations, a plot of the sum of the mean square errors of the recombination fraction estimates verses the sequencing depth is given in Figure 4. This plot suggests that the optimal sequencing depth was around three or four as the mean square error was lowest around these depths.

**Figure 4.**
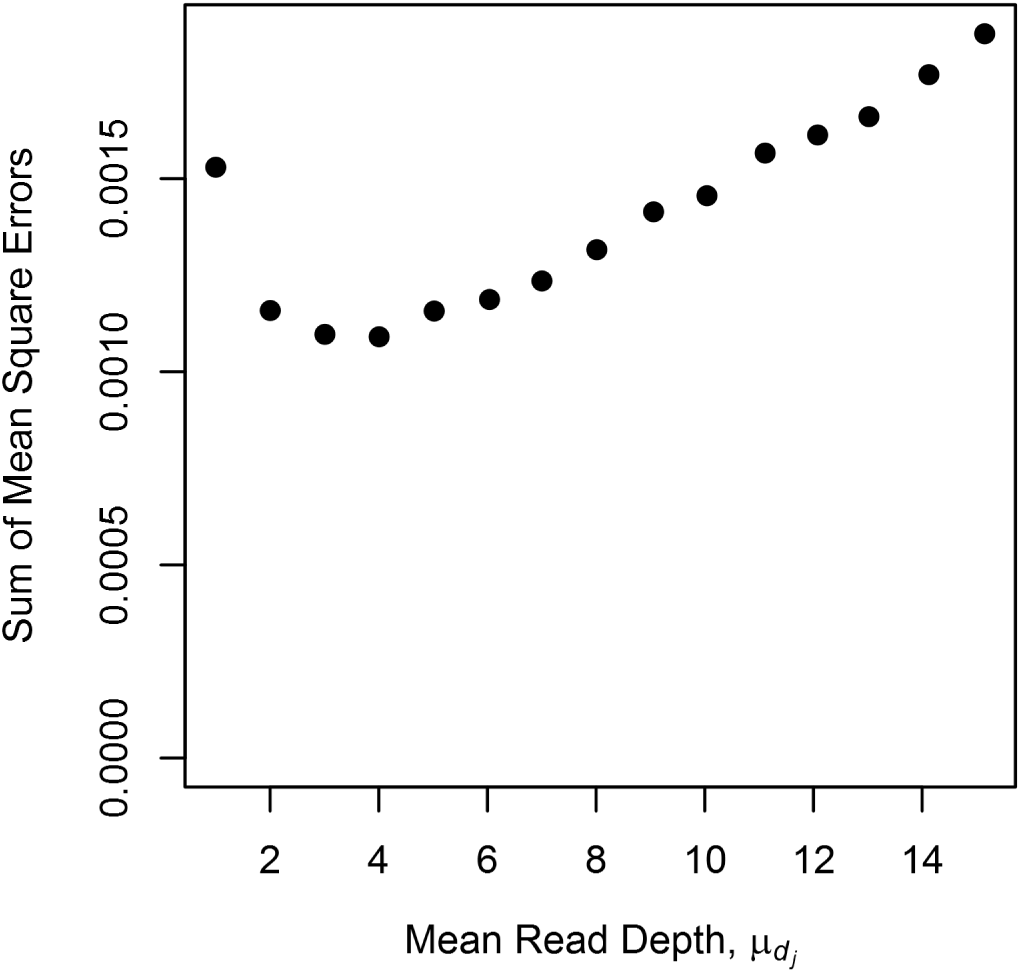
Sum of recombination fraction estimates mean square errors for fixed sequencing effort. Recombination fraction estimates were computed using GM, where the OPGP is known and the sequencing effort was fixed at 10,000 reads. The parameters used to generate the data sets corresponds to the first set of simulations, with the exception that the mean depth and number of individuals were set to maintain a sequencing effort of 10,000. The sum of the mean square errors was calculated using 
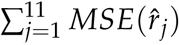
 The number of individuals range from 833 for a mean depth of 1 to 55 for a mean read depth of 15.15.

### Mānuka data

Heatmaps of the 2-point recombination fraction estimates for SNPs located on chromosome 11 are given in Figure S11A in File S1 for SNPs segregating in the paternal parent and in Figure S1B in File S1 for SNPs segregating in the maternal parent. A number of SNPs appeared either to be incorrectly ordered on the chromosome or located on the wrong chromosome and therefore were discarded from the analysis (164 in total). The heatmaps of the remaining SNPs (see Figure S11C and Figure S11D in File S1) suggests that the order of these SNPs was fairly accurate. For the remainder of this analysis, we assume that this order is correct.

Linkage maps of chromosome 11 were computed for both the Low Depth and High Depth set of SNPs using GM and the standard software packages. These linkage maps are given in Figure S12 in File S1 (all maps) and Figure 5 (maps that were less than 150 centimorgans (cM)), with the overall map distance estimates given in Table 5. For the Low Depth set, the maps obtained from LM2, OM, CM and JM were between eight to nine times longer compared to the High Depth set. These inflated map estimates were expected given the substantial proportion of under-called heterozygous genotypes present at low depth and is consistent with the simulation results. Compared to GM and LM2*ε* at high depth, LM2*ε* produced a map that was approximately 20 cM longer when using the Low Depth SNPs, with large distances between the SNPs at the chromosome ends. For the high depth setting, the maps produced by LM2*ε* and GM were similar in length and shorter than the maps obtain using LM2, OM, CM and JM by approximately 30 cM. This suggests that there was sequencing error present in this data set, where both LM2*ε* and GM are accounting for these errors. The overall map distance estimated using GM was consistent across both SNP sets at approximately 76.5 cM, with estimated sequencing error rates of 0.31% for the Low Depth SNPs and 0.20% for the High Depth SNPs. Overall, these results resemble those observed in the simulations and suggests that GM has accounted for most of the errors present in both the low and high depth settings. In terms of phasing, all packages inferred the same phase under both SNP sets, apart from CM which does not require the parental phase to compute the recombination fractions.

**Figure 5.**
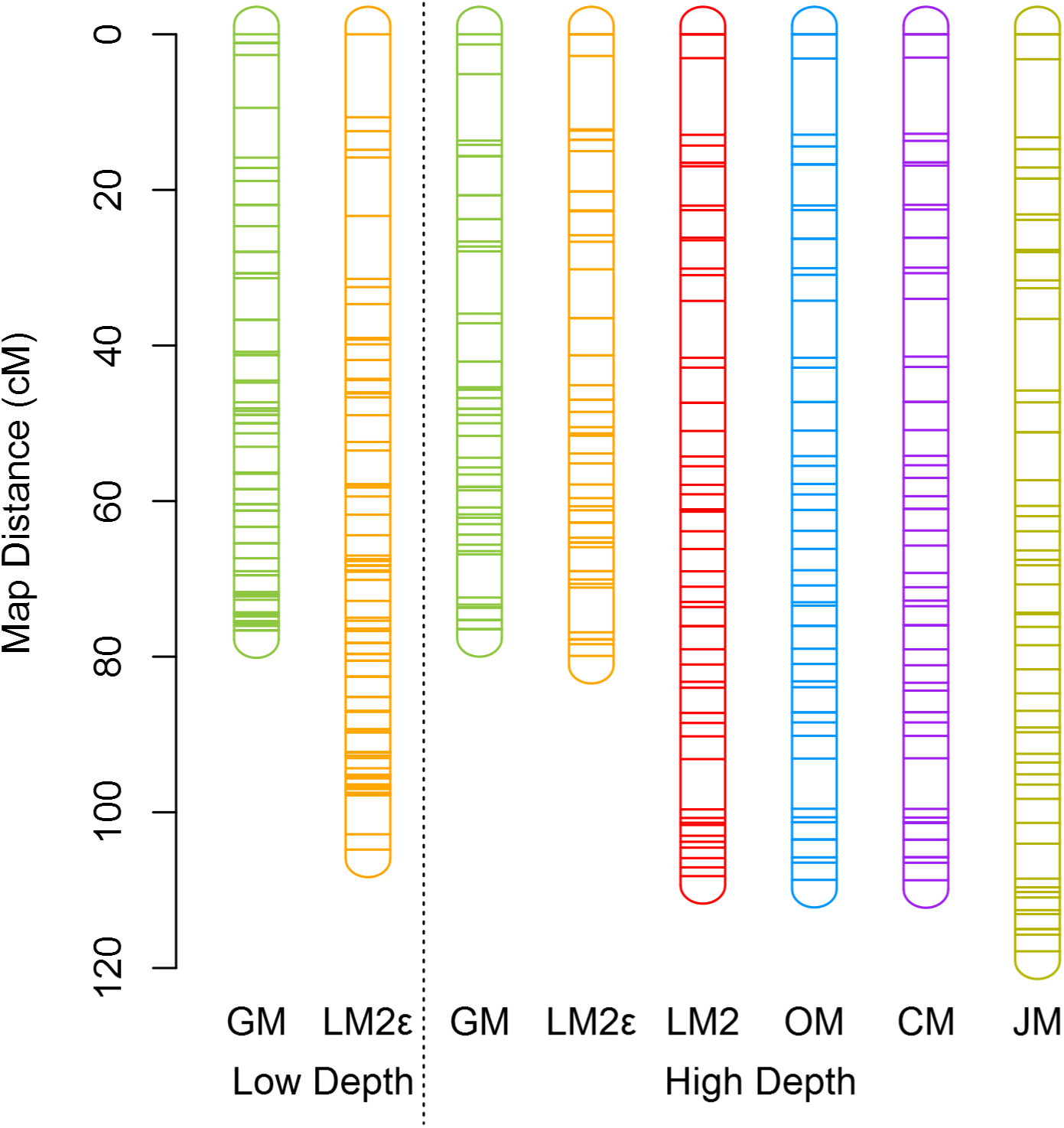
Subset of linkage maps for chromosome 11 of mānuka computed using the various software packages. Low Depth refers to the maps produced using SNPs with a mean read depth below 6, while High Depth refers to maps produced using SNPs with less than 20% missing data after setting genotypes with a read depth below 20 to missing. Map distances are in centimorgans and were computed using the Haldane mapping function.

**Table 5.** Overall map distance estimates (cM) for chromosome 11 of mānuka

| SNP set | GM | LM2ε | LM2 | OM | CM | JM |
| --- | --- | --- | --- | --- | --- | --- |
| Low Depth | 76.6 | 104.8 | 990.0 | 989.1 | 977.1 | 950.7 |
| High Depth | 76.5 | 79.9 | 108.2 | 108.7 | 108.7 | 117.9 |

Map distances were computed using the Haldane mapping function.

## Discussion

We have developed a new statistical method for constructing genetic maps from a set of ordered loci on outcrossed full-sib families in diploid species that have been genotyped using multiplexing sequencing methods. Our methodology uses a HMM approach to overcome the issues associated with mapping in full-sib families and to account for errors resulting from low sequencing depth and miscalled bases. In addition, our methodology is applicable to multi-family and sex-specific situations and has been implemented in the software package GUSMap.

Our simulation results show that GUSMap was able to accurately estimate the recombination fractions and overall map distance for varying sequencing error rates and read mean depth scenarios. In contrast, most of the other software packages were unable to produce reasonable results when there were errors present in the data resulting in biased estimates. In particular, the overall map distances obtained using these packages were inflated, which is known to occur in linkage mapping when genotypes errors are present and not taken into account (Cartwright *et al*. 2007; Lincoln and Lander 1992). Of all the standard software programs, the implementation of Lep-MAP2 that included error parameters was less sensitive to inflation in the overall map distance estimates and was able to provide reasonable results when the number of erroneous genotypes was not too large. However, in the low coverage settings, it still contained substantial bias in the map distance estimates and distortion in the distribution of the SNPs across the map. The maps produced by the packages CRI-MAP, OneMap and Lep-MAP2 without an error parameter gave very similar results since these packages are essentially implementing the same model. In comparison, JoinMap tended to produce maps that were slightly longer in length.

The analysis of the mānuka data set suggests that GUSMap performs well under real life low depth sequencing scenarios. This observation is based on the fact that GUSMap produced consistent estimates of the overall map distance from two independent set of SNPs with different sequencing coverage, and gave a similar map length compared with LM2*ε* at high depth. In contrast, the other packages all produced hugely inflated genetic maps, except for LM2*ε*, although there still appeared to be some inflation under the low depth setting. Thus, this analysis shows that GUSMap is able to reduce map inflation caused by errors in the data and provide better linkage maps and estimates of overall map distance, particularly in low coverage sequencing scenarios.

Of the software packages considered in this paper, only Lep-MAP2 and GUSMap account for errors using information provided by the observed data. For Lep-MAP2, estimation of these errors seems to be based on detecting double recombinants. Thus, the error parameters for the end loci are always zero, since double recombinants cannot be counted, resulting in bias for recombination fraction estimates that include an outside locus when these loci contain erroneous genotype calls. Furthermore, for situations when many errors are present in the data, such as low coverage data, determining which genotypes are incorrect based on double recombinants is difficult. GUSMap, on the other hand, uses the allele count information to model errors due to missing parental alleles and sequencing errors. In particular, it is able to differentiate between the two error types, allowing it to produce accurate maps in low coverage scenarios. What is more, GUSMap can account for errors associated with low read depths in a 2-point analysis, which is not the case when error estimation is based on detecting double recombinants. This is particularly useful for producing heatmaps of 2-point recombination fraction estimates to examine chromosomal ordering. GUSMap also uses only a single parameter to model sequencing error, whereas LM2*ε* specifies a separate error parameter for each locus. Consequently, GUSMap makes the assumption that the sequencing error rate is constant across individuals and loci. In practice, this assumption may not hold, although the mānuka results suggest that it may be reasonable in some situations.

The simulation results suggest that GUSMap is able to provide reasonably accurate estimates of the sequencing error rate, although there was a small bias at low depth. For the mānuka data set, the estimates from GUSMap suggest that between 0.2% to 0.3% of the reads in the filtered data were sequencing errors, although there was discrepancy in the sequencing error rates estimated between the two SNP sets. This discrepancy could be due to natural variation, particularly as the simulation results suggest that there is large variability in the sequencing error estimates at low depth. Alternatively, it could be that high depth SNPs with large sequencing errors were removed through the filtering process as detection of these SNPs would be easier since the sequencing errors are not confounded with the errors resulting from low sequencing depth.

Nearly all full-sib family software packages require inferring parental phase. Phasing errors can result in estimates that are close to or equal to 0.5. GUSMap and Lep-MAP2 were mostly able to correctly infer phase across all the simulations. Both of these packages use a similar phasing approach in that they infer phase based on sex-specific recombination fractions estimated on the interval [0, 1] using a multipoint likelihood containing all the loci. In contrast, OneMap failed to correctly infer phase in the low depth simulations, which suggests that phase inference based on maximizing the likelihood value is unreliable in the presence of severe model misspecification. JoinMap also failed to infer phase in some data sets, which suggests that phasing using a multipoint approach can be superior to using a 3-point approach. The ability to correctly infer phase also depends on a number of factors; namely, the density of the markers, the family size, and for sequencing data, the average sequencing depth. Simulation results (Figure S13 in File S1) suggest that GUSMap is able to infer phase for low coverage data provided that the maps are at moderate-to-high density, the mean read depth is at least 2, and that there are at least 25 progeny in the family.

A number of assumptions have been made in the methodology we have outlined. Firstly, the order of the loci is assumed to be known beforehand, which is often not the case with sequencing data, particularly for *de novo* assemblies. One approach to ordering loci is to evaluate the likelihood under different chromosomal orders, where the best order is the one that gives the highest likelihood value. This approach is feasible for improving order locally, provided that the initial order is fairly accurate, but is impractical for ordering large number of loci that are randomly ordered. A reasonable initial order could be computed by combining 2-point estimates obtained from GUSMap with existing ordering algorithms. More research is required to investigate loci ordering in low coverage settings. Another assumption is that all the parental genotypes are known for all loci, so that the STs can be determined unequivocally. In practice, all the parents will be sequenced using multiplexing methods and therefore are subject to the same type of genotyping errors as the progeny. One way to circumvent this issue is to sequence the parents multiple times to obtain higher sequencing depths, although this still results in some loci having insufficient depths to accurately infer the ST. Alternatively, if there is a sufficient number of individuals in each family, the ST of each locus could be inferred from progeny genotypes using a segregation test. Other assumptions include independence of the reads observed in the sequencing data between loci, which is a reasonable assumption provided there is only a single locus on each read, and the sampling of the alleles from the true genotype is random. For the latter assumption, the probabilities in Eq (9) can be adjusted to reflect any prior knowledge of the sampling of the true genotypes (e.g., preferential sampling of alleles). This methodology is limited to autosomal biallelic loci in diploid species or functionally diploid species (e.g., allopolyploids). Extension of this methodology to allosomal (sex-linked) loci and multiallelic loci (e.g., microsatellite markers) would require deriving the correct emission probabilities in Eq (9) for the HMM.

GUSMap provides researchers with a tool to compute genetic maps from a set of ordered loci using sequencing data and overcomes a number of issues related to this data. Firstly, it is able to handle varying sequencing depths across SNPs, which is typical of sequencing data, allowing more SNPs to be utilized that would otherwise be discarded in a high depth analysis. Secondly, SNPs called in the bioinformatics process must meet a minimal set of filtering criteria, which is aimed at removing erroneous genotypes and fictitious SNPs. GUSMap removes the need for filtering based on discarding genotyping calls based on read depth information or skewed apparent segregation and correcting erroneous genotypes through detecting double recombinants. This allows researchers to use low coverage data, especially when cost constraints may prohibit the production of sufficiently high coverage data, to construct genetic maps. Another advantage of GUSMap is its use of a statistical approach to model errors, which allows it to be combined with existing statistical techniques to make inference on model parameters, such as quantifying the rate of sequencing errors, and assessing modeling assumptions. Although the methodology of GUSMap is derived specifically for outbred full-sib populations, it can also be applied to inbred backcross populations, since the segregation type of all the loci are either paternal-informative or maternal-informative, and inbred F2 populations where the segregation type of all the loci are both-informative.

## Acknowledgements

This work was funded by a University of Otago Doctoral Scholarship to TPB and by the Ministry of Business, Innovation and Employment (New Zealand) via its funding of the "Genomics for Production & Security in a Biological Economy" programme (Contract C10X1306) to AgResearch Ltd and via its SSIF funding "Discovery Science" to Plant & Food Research. The mānuka plants originated from the New Zealand East Cape region and used as parents for a control cross were obtained in collaboration with Horouta Manuka Company Ltd and the use of the seedlings for genetic mapping was approved by Ngati Porou Miere Ltd. We thank Dr David Lewis and Mrs Julie Ryan for making the cross and raising the seedlings, Mrs Tracey Van Stijn for the sequencing of the mānuka samples, Dr Rudiger Brauning for bioinformatics support and Mr Maxim Mikhisor for assistance with programming the simulations.

